# IQMMA: an efficient MS1 intensity extraction using multiple feature detection algorithms for DDA proteomics

**DOI:** 10.1101/2023.02.03.526776

**Authors:** Valeriy I. Postoenko, Leyla A. Garibova, Lev I. Levitsky, Julia A. Bubis, Mikhail V. Gorshkov, Mark V. Ivanov

**Author notes:** These authors contributed equally (Valeriy I. Postoenko, Leyla A. Garibova).

## Abstract

One of the key steps in data dependent acquisition (DDA) proteomics is detection of peptide isotopic clusters, also called «features», in MS1 spectra and matching them to MS/MS-based peptide identifications. A number of peptide feature detection tools became available in recent years each relying on its own matching algorithm. Here, we provide an integrated solution, Intensity-based Quantitative Mix and Match Approach (IQMMA), which integrates a number of peptide feature detection algorithms and returns the most probable intensity values for the MS/MS-based identifications. IQMMA was tested using available proteomic data acquired for both well-characterized (ground truth) and real-world biological samples, including a mix of *E. Coli* digest spiked at different concentrations into the HeLa digest used as a background and a set of glioblastoma cell lines. Three open-source feature detection algorithms were integrated: Dinosaur, biosaur2 and OpenMS FeatureFinder. Neither of them was found optimal when applied individually to all the datasets employed in this work, however, their combined use in IQMMA improved efficiency of subsequent protein quantitation. The software implementing IQMMA is freely available at https://github.com/PostoenkoVI/IQMMA under Apache 2.0 license.

## Introduction

Currently, many applications in biomedical research become heavily relying on quantitative proteome analyses^1,2^. Mass spectrometry (MS) in combination with liquid chromatography is the method of choice in these analyses allowing proteome-wide protein identification and quantitation. Two main approaches are used for MS-based proteome-wide analysis: data-independent^3^ (DIA) and data-dependent^4^ (DDA) acquisition, both based on the so-called bottom-up approach, in which the proteins are first enzymatically digested followed by on-line separation of proteolytic mixture at the front-end of a mass spectrometer and identification of peptides from their fragmentation mass spectra (MS/MS). Protein quantitation is one of the final steps of this proteomic workflow albeit a crucial one. Quantitative analysis is performed based on stable isotope labeling or using label-free approaches. The former requires extra sample preparation and has additional issues from co-isolated and co-fragmented ions when it comes to the so-called chimeric spectra^5,6^. Label-free quantitation has no such limitations but strongly relies on data processing algorithms. In particular, the DIA methods utilize peptide ion intensities in MS/MS spectra, and the main source of error here is due to the summation of fragment ion intensities of co-fragmented precursor ions. Quantitation in DDA methods is usually done by spectral counting algorithms such as NSAF^7^, emPAI^8^ or SI_N_^9^. These methods rely on the number of identified PSMs per protein and show low consistency due to the stochastic nature of precursor selection and dynamic exclusion processes^10^. Intensities of MS/MS spectra in the DDA approach also do not provide accurate quantitative information about peptides, as MS/MS spectra are produced with uncertainty within the peptide elution profile and cannot be directly compared between experimental runs. Alternatively, one of the widely explored approaches for accurate label-free quantitation is extraction of MS1 peptide intensities for identified MS/MS spectra^11,12^. The key step in the extraction of intensities is detection of the so-called peptide-like features in the MS1 spectra^13^ representing the peptide ^13^C-isotopic distribution (cluster). Note that the number of peptide ions detected in each MS1 spectrum is typically high for complex samples, and a significant number of peaks cannot be unambiguously interpreted as peptide features. A range of peptide feature detection methods exist, implementing different algorithms and providing different sets of detected isotopic clusters. These detected features should be mapped to identified MS/MS spectra using *m/z*, retention time, charge and ion mobility information^14,15^. Different implementations exist for such mapping, including those built into proteome analysis pipelines (MaxQuant^16,17^, FragPipe^18^), feature detection (OpenMS^19^) or quantitation (Triqler^20^). There are multiple reasons why this mapping cannot be resolved unambiguously: different sets of peptide features generated using different feature detection algorithms, multiple feature candidates per MS/MS spectrum, missing features, etc. However, the idea of combining multiple algorithms to obtain optimal results is not uncommon in proteomics. For example, the SearchGUI^21^ software uses multiple search engines to efficiently identify peptides. It is well-known that intersection of peptide identifications between multiple search engines rarely exceeds 80%^22^, which leaves room for improvement. A similar approach is conceivable for MS1 feature detection and quantitative analysis, however, it was barely explored. A number of recent studies aiming at benchmarking the protein quantitation were focused mostly on the entire quantitation workflows, rather than on the specifics of the peptide feature detection methods^23^. While some previous reports have demonstrated integration of quantitation methods, such as combining label-free and labeled approaches^24^, or combination of the quantitation results obtained for different datasets^25^, there is a lack of implementations of the integration of different standalone quantitation workflows at the peptide feature detection level. In this study we analyzed the differences in quantitation results between feature detection algorithms combined with a particular quantitation method. Employment of multiple feature detection algorithms in a single quantitation analysis is proposed and implemented as a standalone, versatile software tool called IQMMA (Intensity-based Quantitative Mix and Match Approach).

## Experimental section

### HeLa-*E. coli* samples

LC-MS/MS data were used from a previous study^26^ (ProteomeXchange id PXD001385). Briefly, *E. coli* cell lysate in concentrations of 3, 7.5, 10 and 15 ng was spiked into 200 ng of HeLa cell lysate. LC-MS/MS analysis in DDA mode was performed using Q Exactive Plus mass spectrometer (Thermo Scientific, San Jose, CA, USA) coupled with nano-ACQUITY UPLC system (Waters, Milford, MA, USA) and 105 min LC gradient.

### Glioblastoma multiforme (GBM) samples

LC-MS/MS experiments with low-passage culture of GBM (3821) cell lines were used from a previous study^27^ (ProteomeXchange id PXD022906). Briefly, cell cultures were grown in four replicates and treated with recombinant interferon (IFN) α-2b (4 control, 4 treated samples, 2 technical repeats each). LC-MS/MS analysis was performed in DDA mode using Q Exactive HF mass spectrometer (Thermo Scientific, Waltham, MA, USA) coupled with an UltiMate 3000 nanoflow LC system (Thermo Scientific, Bremen, Germany) and 120 min LC gradient.

#### Data Analysis

Raw files were converted into mzML format using MSConvert^28^ (v. 3.0.20287).

Peptide features were detected using biosaur2^14^ (v. 0.2.3), Dinosaur^29^ (v. 1.2.0) and OpenMS FeatureFinderCentroided^19^ (v. 2.7.0).

Peptide identification was done using IdentiPy^30^ search engine. The parameters for search were as follows: 10 ppm precursor mass accuracy, 0.05 Da fragment mass accuracy, 2 missed cleavages, carbamidomethylation of cysteine as fixed modification, minimal peptide length of six amino acids. Possible C13 isotope errors of one or two units were allowed. Postsearch analysis was performed using Scavager^31^ (v. 0.2.9). Diffacto^32^ (v. 1.0.6) was used for protein quantitation with a global normalization strategy on median values.

## Results and Discussion

### Workflow

A general scheme of the workflow is shown in Figure 1. IQMMA requires mzML files and PSM lists produced by database search and, optionally, postsearch validation. At the first step, the programs for feature detection (Dinosaur, biosaur2, OpenMS FeatureFinderCentroided) are sequentially launched to generate three sets of MS1 peptide isotopic cluster features for each analyzed mzML file. Each set of features is then mapped to the identified MS/MS-based peptide-spectrum matches (PSMs) to produce peptide-feature pairs that are used in further analysis. The mapping is done by default with no match-between-runs (MBR) functionality which is integrated within the IQMMA software and can be optionally turned on. Details on the mapping procedure are discussed below. Next, for each of the three sets of PSMs with mapped MS1 intensities, intermediate Diffacto analyses evaluate the results of quantitation performed separately for each feature detection algorithm. This is followed by the procedure of mixing the feature detection algorithms applied for each PSM. This procedure determines the most appropriate sets of intensities of the three feature detection methods (Figure 1B) according to two criteria: first, the set with the lowest number of missing values is selected; if several sets have the lowest number of missing values, then the set with the lowest sum of the squared normalized coefficients of variation (CV) for intensities in two samples is chosen. Note, all CV values are normalized by the feature detection median CV value to eliminate systematic biases. Finally, the mixed set of intensities for PSMs is used by Diffacto to obtain the final results of quantitation.

**Figure 1.**
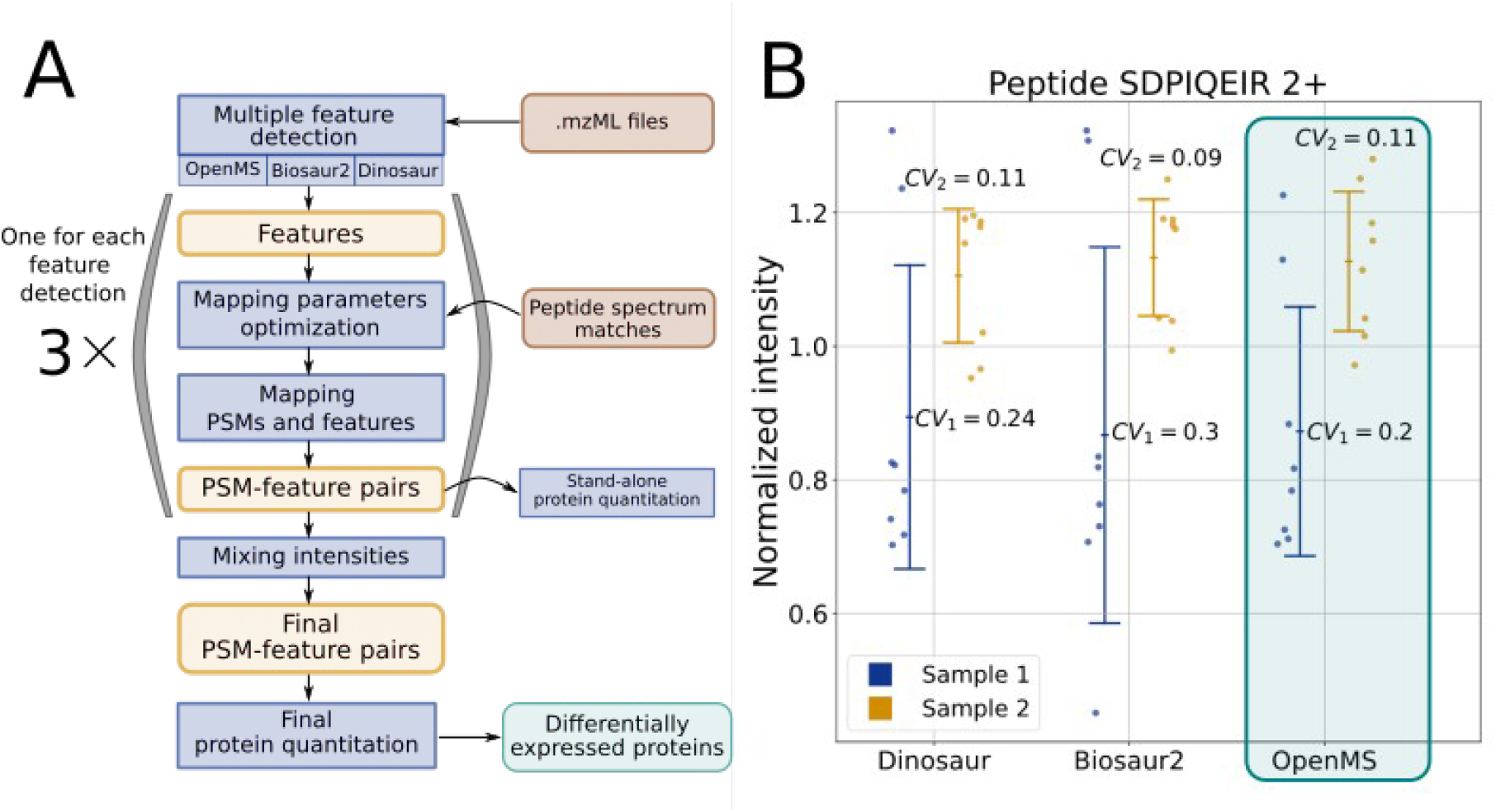
(A) General IQMMA workflow scheme. (B) Choosing the set of intensities for an individual peptide by minimizing the sum of squares of CV_1_ and CV_2_ - coefficients of variations for intensities of the peptide in two quantitation sample groups.

### Feature mapping algorithm

#### Accounting for mass differences

First, a preliminary matching of a peptide mass in MS/MS spectra to the one in MS1 spectra is performed using 100 ppm mass tolerance. The distribution of mass differences between MS/MS-based precursor m/z and MS1 peptide ion m/z is expected to be a sum of normal (true matches) and uniform (false matches) distributions. IQMMA fits experimentally obtained distribution to this sum and estimates mean and standard deviation for the normal distribution (Fig. 2A). The mean +/- 6 standard deviations (determined empirically in this work based on the well-characterized dataset) is used as the mass difference threshold in the final match of the feature between MS1 and MS/MS spectra.

**Figure 2.**
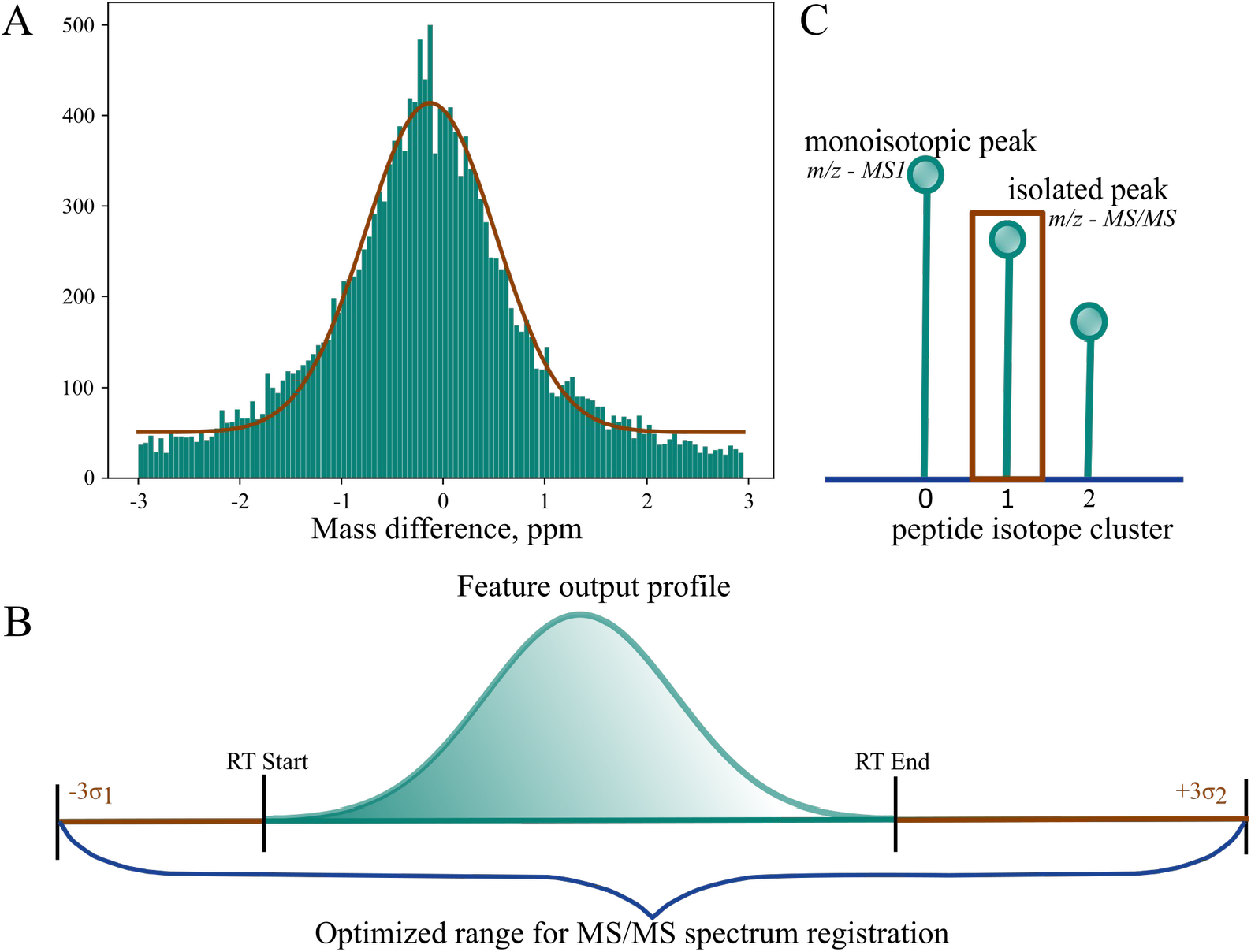
Optimization procedures used in IQMMA. (A) Mass difference between MS/MS precursor *m/z* and MS1 peptide feature *m/z* values are fitted using a sum of normal and uniform distributions. Histogram is shown as an example for HeLa - 15 ng *E. coli* single file. (B) Distribution of allowed range for MS/MS spectrum retention time goes beyond the RT profile of MS1 peptide feature. (C) Cases for the errors in peptide isotope cluster selection. Identified MS/MS spectra are mapped to MS1 peptide isotopic clusters using measured *m/z*, retention time (RT), charge, and ion mobility values. The latter are binned by the compensation voltages in case of FAIMS. While charge and FAIMS compensated voltage annotations can be matched directly, *m/z* and RTs differ between MS1 and MS/MS spectra, thus, should be matched as described below.

#### Accounting for RT differences

Typically, MS/MS spectrum acquisition time is usually inside of MS1 peptide feature retention profile. IQMMA additionally takes into account cases when MS/MS spectra are acquired outside of MS1 retention profile, which may happen due to multiple reasons. For example, inaccuracies in MS1 feature detection algorithms may lead to truncation of detected MS1 chromatographic peaks. Another possible source of RT discrepancies are chimeric MS/MS spectra, when peptides were isolated and fragmented without the evidence at MS1 level. Similar to the mass differences, we estimate both mean and standard deviation for the RT difference distribution. However, we analyze separately two RT differences: first one is between MS/MS scan time and MS1 feature starting time, and the second one is between MS/MS scan time and MS1 feature ending time. The optimized allowed range for MS/MS acquisition time is extended outside the MS1 retention profile by 3 estimated standard deviations as shown in Fig. 2B.

#### Accounting for isotopic mass errors

Similar to the isotopic mass error option implemented in MS/MS search engines, we added such an option for MS1 feature mapping. These types of errors may occur while determining the monoisotopic peaks by feature detection algorithms or by acquisition software^33^. Thus, there can be a difference in masses between the MS1 peptide feature and the recorded precursor up to a few Da due to the presence of ^13^C isotopes. One possible outcome from this error is that a wrong *m/z* peak from another peptide ion is annotated as a monoisotopic peak by a feature detection algorithm. At the same time, the mass spectrometer software may correctly select the monoisotopic peak for isolation and fragmentation. In the example shown in Fig. 2C, there will be a -1.00335 Da difference between MS1 and MS/MS precursor masses. By default, the algorithm checks up to five ^13^C isotope related mass errors of -2, -1, 0, +1 and +2. In some cases, multiple MS1 peptide isotopic clusters are matched to the MS/MS-based peptide identification, even after the refinement of *m/z* and RT tolerances. The priority is then given to the matches with minimal absolute isotope mass error. If an ambiguity still remains, the minimum root squared error for mass and RT difference is chosen.

#### Match-between-runs

Match-between-runs (MBR) functionality is implemented in the IQMMA workflow and can be turned on optionally. Once it is turned on, the mapping algorithm is applied for all MS/MS matches and MS1 features from multiple data files. However, the priority in intensity matching is given to the MS1 features mapped to the peptide from the same experimental file.

#### Missing values

“Missing values” in the PSM-feature mapping have two possible reasons: no MS1 feature for an identified MS/MS spectrum, or an MS/MS peptide identifications not reproduced between replicates. MBR functionality is aimed at solving the latter. By combining multiple feature detection algorithms and optimizing the RT ranges for PSM-feature mapping, IQMMA partially solves the first part of the “missing values” problem. 95% of identified PSMs were mapped to any MS1 feature using IQMMA for the HeLa-*E. coli* samples. For the standalone usage of feature detections, the 86%, 90% and 65% of PSMs were mapped to MS1 features for biosaur2, Dinosaur and OpenMS.

### Quantitation benchmark

We kept fixed all the stages of the quantitation workflow except the feature detection and performed six pairwise concentration comparisons of the *HeLa - E.coli* dataset (Figure 3). As an example, the results of comparison obtained for the samples of 15 and 7.5 ng *E. coli* proteins spiked into 200 ng HeLa lysate are shown in Fig.3A as a Venn diagram. Applying IQMMA increases the number of differentially expressed proteins and many of them are unique for the approach. However, some of the *E. coli* proteins reported by the standalone approaches are absent in IQMMA results, which leaves room for further improvements. Fig.3B shows the number of differentially expressed proteins reported by different quantitation workflows. Most of these proteins are *E. coli*, which are true positives, and IQMMA performed the best in all pairwise comparisons. There is no clear second best standalone feature detection approach. This is an interesting observation because all data and samples were obtained within one experimental study. Thus, even preliminary experiments with known samples cannot help with selection of the optimal standalone single feature detection approach. However, the proposed IQMMA algorithm provides both versatility and maximal efficiency. The reported differentially expressed human proteins were considered as false positives and shown with numbers above the bars in Fig. 3B. These numbers are useful for a rough estimation of empirical FDR behind the particular approach. The maximal fraction of human proteins is 6.7% for OpenMS results obtained from “10 ng to 3 ng” comparison. Note that quantitative FDR thresholds were set at 5% (p-values less than 0.05 with Bonferroni multiple comparison correction). Thus, all pairwise comparisons show quite reliable results in most cases and, importantly, IQMMA does not increase the FDR. The volcano plots for all feature detection methods and all pairwise concentration comparisons are shown in Supplementary Figure S1. The quantitation accuracy for a particular approach was estimated then as the root mean squared percentage errors (RMSPE) calculated using the following equation:

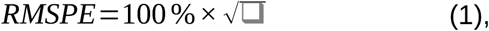

in which N is the number of differentially expressed *E. coli* proteins, *FC*_real_ and *FC*_estimated_ are experimental and reported fold changes in log_2_ scale, respectively. The RMSPE values calculated using Eq.1 are shown in Fig.3C. These calculations were performed only for the intersection of differentially expressed proteins between all 4 approaches and the number of these proteins are shown above the bars in the figure.

**Figure 3.**
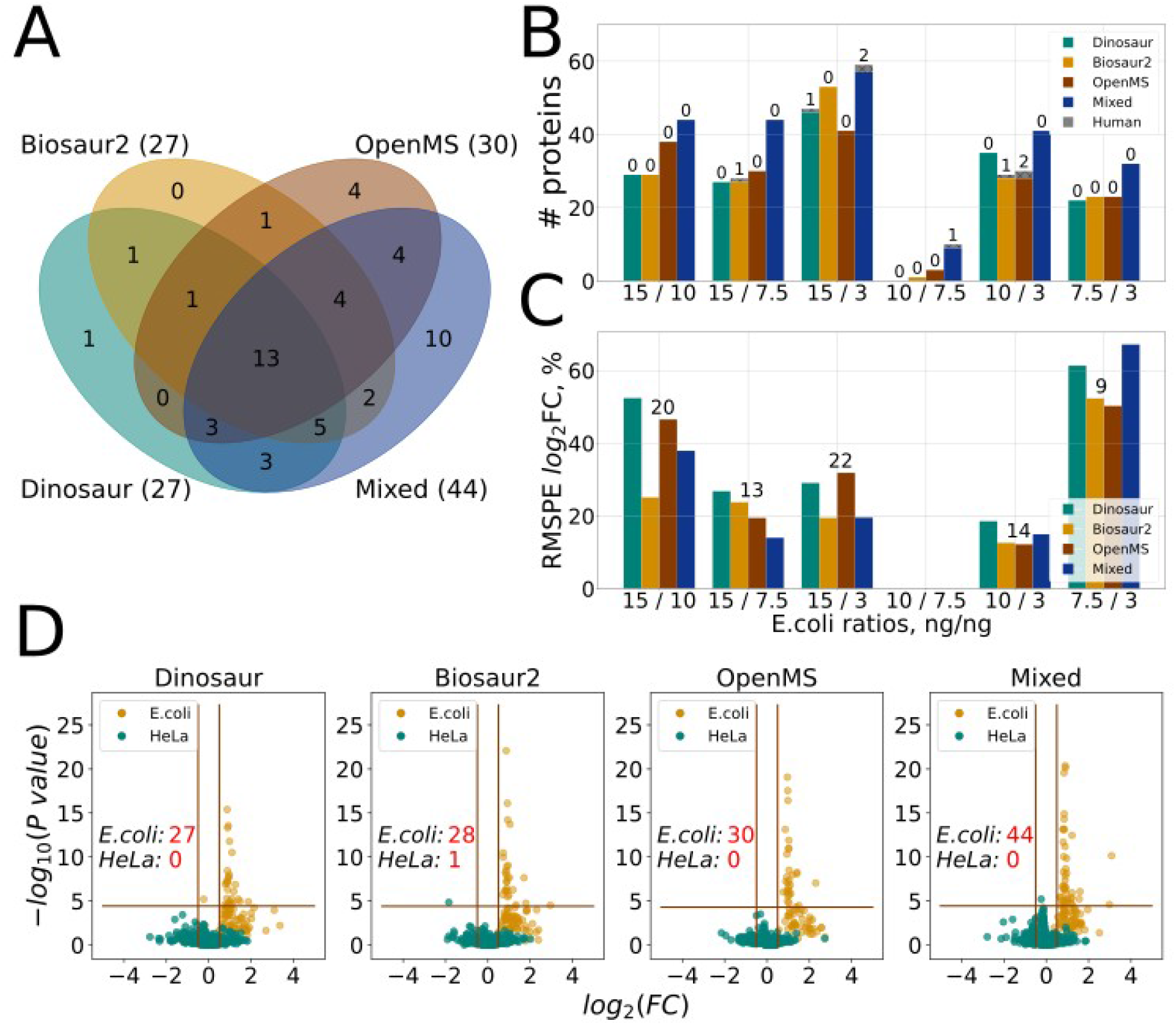
Results of the quantitation analysis of *HeLa* - *E. coli* datasets using IQMMA with three standalone feature detections: Dinosaur, biosaur2 and OpenMS FeatureFinderCentroided: (A) Venn diagram for *E. coli* proteins differentially expressed between the samples with 15 and 7.5 ng of *E. coli* lysate spiked into 200 ng Hela lysate; (B and C) histograms for six pairwise concentration comparisons for all Hela-*E. coli* datasets: total detected differentially expressed proteins (B) and rooted mean squared percentage error of log2FC (C). The numbers above the bars show the quantity of human proteins falsely identified as differentially expressed (B), and the size of the set of the common proteins for all four methods, for which the error was calculated (C); (D) example of volcano plots used to identify differentially expressed proteins between the samples corresponding for spiking *E. coli* proteins at amount of 15 and 7.5 ng. Horizontal line represents 0.05 p-value threshold adjusted using Bonferroni correction, vertical lines represent fold change thresholds, which were set at 0.5 as a half of the real E. coli proteins ratio.

The isotope assignment error problem was described above in the “feature mapping algorithm” section. Accounting this error for matching MS1 peptide features and identified MS/MS spectra increases the number of candidates for matching, resulting in the increase in both false and true MS1-MS/MS matches. There is no way to evaluate the real data apriori on whether additional matches are true or false. However, the quantitation results can provide an indirect assessment for such matches. The average number of reported differentially expressed *E. coli* proteins for different isotope assignment errors were 41, 44.6, 44.2, 44.8 and 47 for [0], [0, 1], [0, 1, -1], [0, 1, -1, 2] and [0, 1, -1, 2, -2] sets of isotopic errors, respectively. The average number of reported differentially expressed human proteins were 0.2, 0.4, 0.4, 0.4 and 0.4 for the same sets, respectively. These numbers were calculated as an average for all 6 pairwise comparisons of concentrations. Accounting for the isotope assignment errors increases the number of reported *E. coli* proteins which means that such procedure adds more benefit from true MS1-MS/MS matches than harm from false ones.

To test the IQMMA algorithm further, the Glioblastoma dataset was taken as an example of real-world quantitative proteomics study^27^. The proposed algorithm provided more differentially expressed proteins compared to the standalone application of feature detections (see supplementary figure S2). The numbers of differentially expressed proteins were 27, 23, 25 and 28 for the Dinosaur, biosaur2, OpenMS and IQMMA results, respectively. The gene expression analysis of these proteins has revealed 8, 10, 9 and 10 genes from the type I interferon signaling pathway for the Dinosaur, biosaur2, OpenMS and IQMMA results, respectively.

All quantitations were performed in the manuscript with turned off match-between-runs option. The HeLa-*E. coli* results using MBR are shown in Supplementary figures S3 and S4. MBR increases the number of differentially expressed *E. coli* proteins for all feature detection methods. However, this option increases quantitative empirical FDR calculated using differentially expressed human proteins as well. The latter agrees with the recent Lim et al work^34^. Our results show that the proposed IQMMA method works also for the MBR approach and does not increase FDR higher than in the standalone feature detection method. Also, MBR provides more accurate Fold change estimation for our data (see Supplementary Figure S4B and Figure 3C). Nevertheless, the key focus of our work is to show that the proposed IQMMA approach is universal for both cases, rather than characterization of MBR.

### Software details

Using multiple feature detection tools instead of a single one entails an increase in processing time. We performed a speed comparison of each stage of the workflow for the IQMMA using a computer with 6 cores Intel(R) Core(TM) i7-3930K CPU (see Fig. 4). The 6 raw files of the HeLa-*E. coli* data contain 169 515 MS/MS spectra and 10.5 hours of LC-MS acquisition time in total. According to the test, most of the processing time is spent by feature detection in general and by the OpenMS FeatureFinderCentroided, in particular. The IQMMA software has an ability to reuse the previously generated features and “skip” the feature detection in case of data reanalysis, which can reduce 20-fold the re-analysis time.

**Figure 4.**
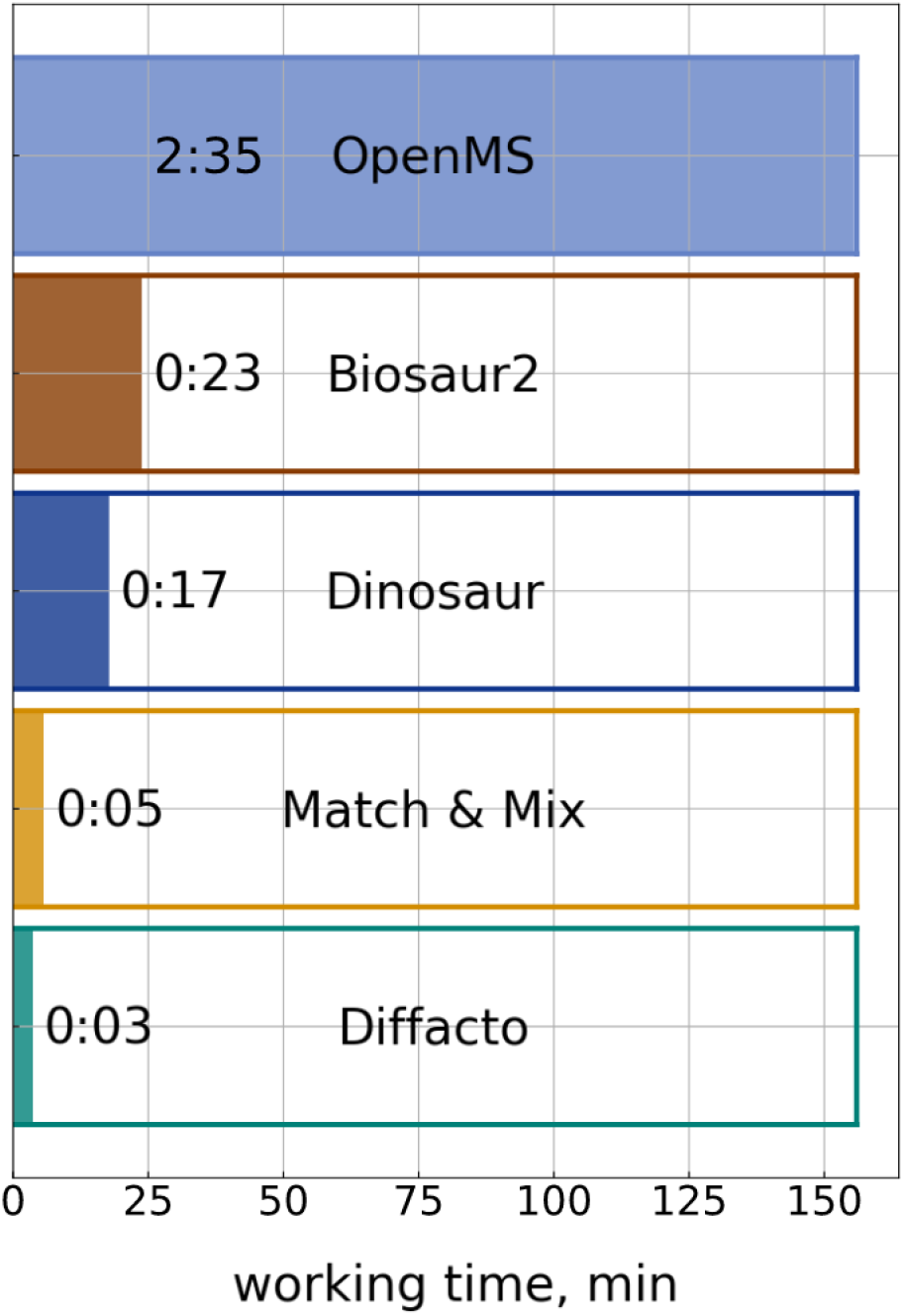
Working time diagram showing distribution of processing time between all stages. Human - *E.coli* data set, 15 to 7.5 ng of *E. coli* proteins.

There are two possible ways of using IQMMA. The first one is a full quantitation pipeline for two samples, starting from the identified peptides and ending with quantitation results. The second one is a simple mapping of MS1 features from a single feature detection result to MS/MS identifications in a single identification results file. The latter can be used by researchers willing to proceed with quantitation analysis manually or by using a quantitation algorithm other than Diffacto. However, the full pipeline is needed to activate the Mixing algorithm, since it depends on analysis of multiple replicates and samples.

IQMMA supports peptide identification results in pepXML and mzIdentML formats, making it compatible with most search engines, including MSFragger^35^, MS-GF+^36^ and IdentiPy^30^. It also supports tab-separated files with predefined column names, making it compatible with most available pipelines and methods. However, the best way to use a full pipeline is the one with added postsearch validation. The Scavager postsearch algorithm is supported out of the box, which itself supports most of the proteomics search engines.

## Conclusions

We developed a method and software for mapping MS/MS peptide identification to MS1 peptide isotopic clusters. The method utilizes multiple feature detection algorithms, which results in more sensitive protein quantitation with an average of 49% increase in the number of differentially expressed proteins. Specifically, we have demonstrated application of the proposed method on well-characterized (ground truth) and biologically relevant samples. Software is freely available and can be installed at https://github.com/PostoenkoVI/IQMMA under Apache 2.0 license.

## Supporting Information

**1. Supporting figure S1.** Volcano plots comparison for Human - *E. coli* dataset with turned off match-between-runs option.

**2. Supporting figure S2.** Volcano plots comparison for glioblastoma dataset.

**3. Supporting figure S3.** Volcano plots comparison for Human - *E. coli* dataset with turned on match-between-runs option.

**4. Supporting figure S4.** Results of the quantitation analysis using the IQMMA approach for Human - *E. coli* dataset with turned on match-between-runs option.

## ACKNOWLEDGMENTS

The work was supported by the Russian Science Foundation, grant no. 21-74-10128 to M.V.I.

## For Table of Contents Only

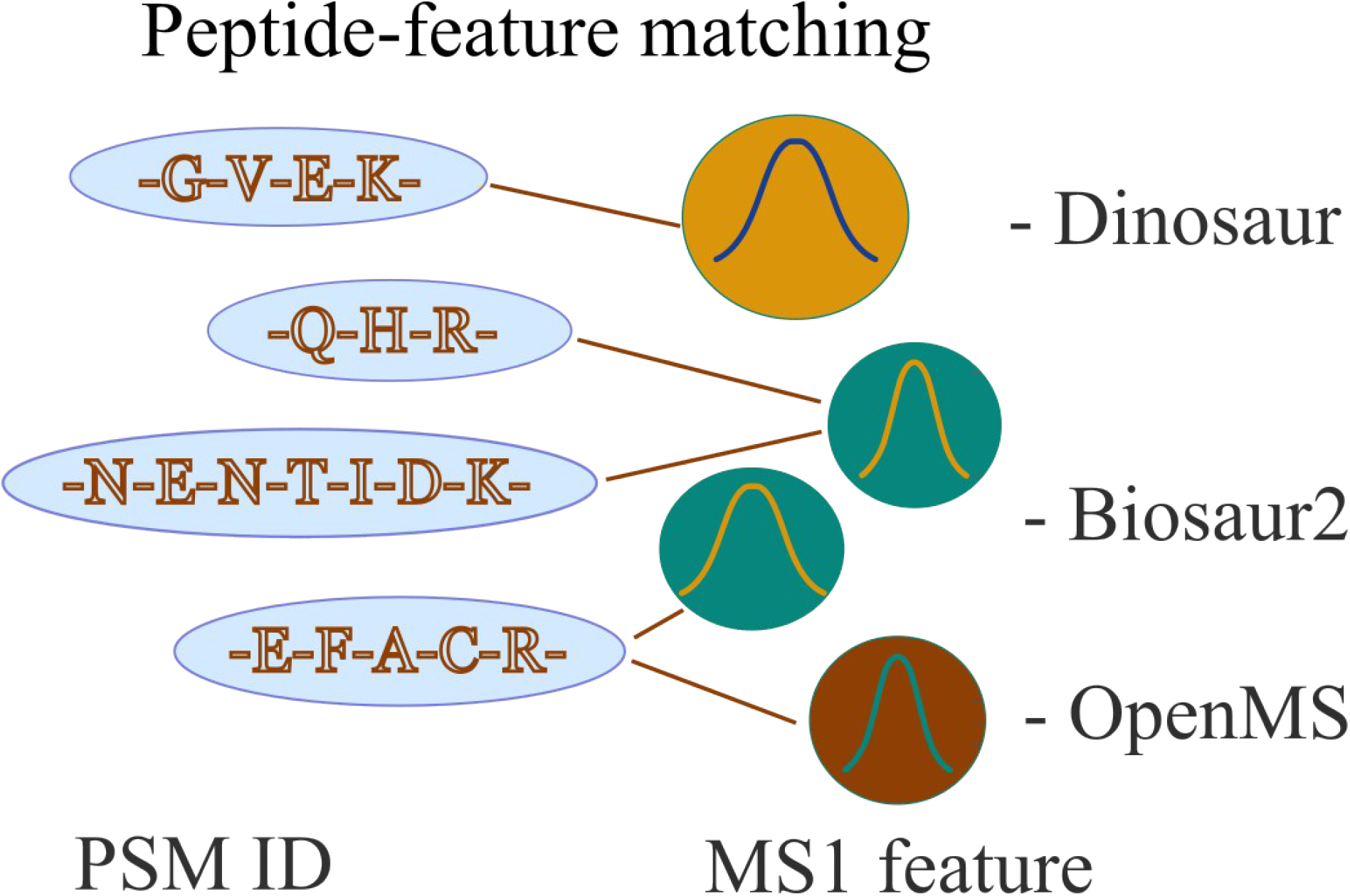

