## Supporting Information for "IQMMA: an efficient MS1 intensity extraction using multiple feature detection algorithms for DDA proteomics"

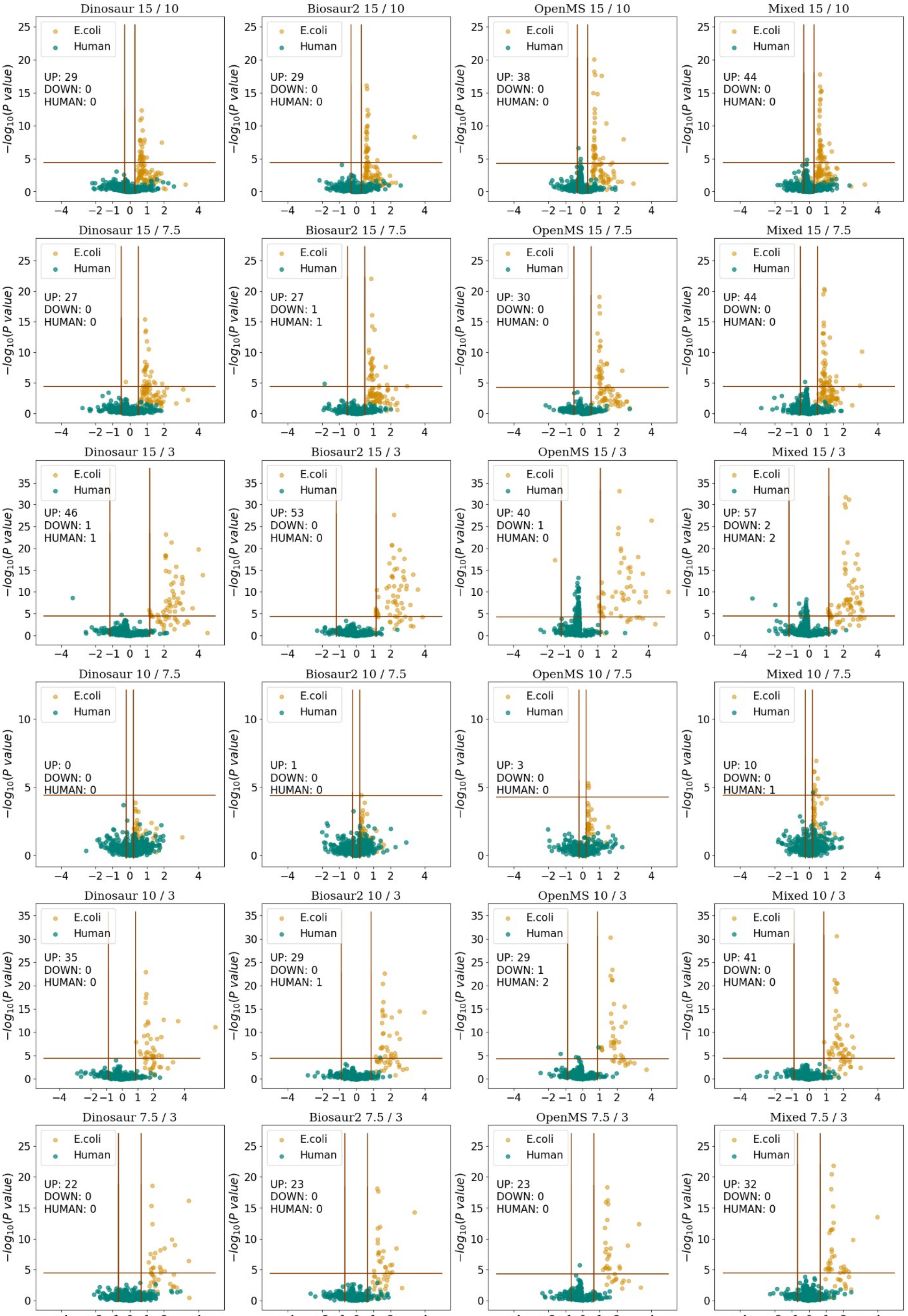


**Supplementary figure S1.** Volcano plots comparison for Human - *E. coli* dataset with turned off match-between-runs option. Horizontal line represents 0.05 p-value threshold adjusted using Bonferroni correction, vertical lines represent fold change thresholds which were set at a half of the real *E. coli* proteins ratio.

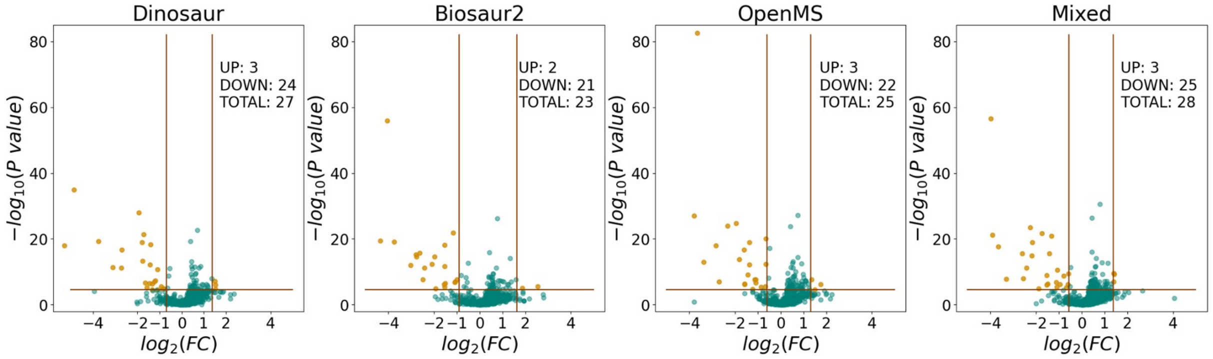


**Supplementary figure S2.** Quantitation results (volcano plots) for glioblastoma data set. Horizontal line represents 0.05 p-value threshold adjusted using Bonferroni correction, vertical lines represent fold change thresholds which were set according to estimation of distribution of proteins under p-value threshold.

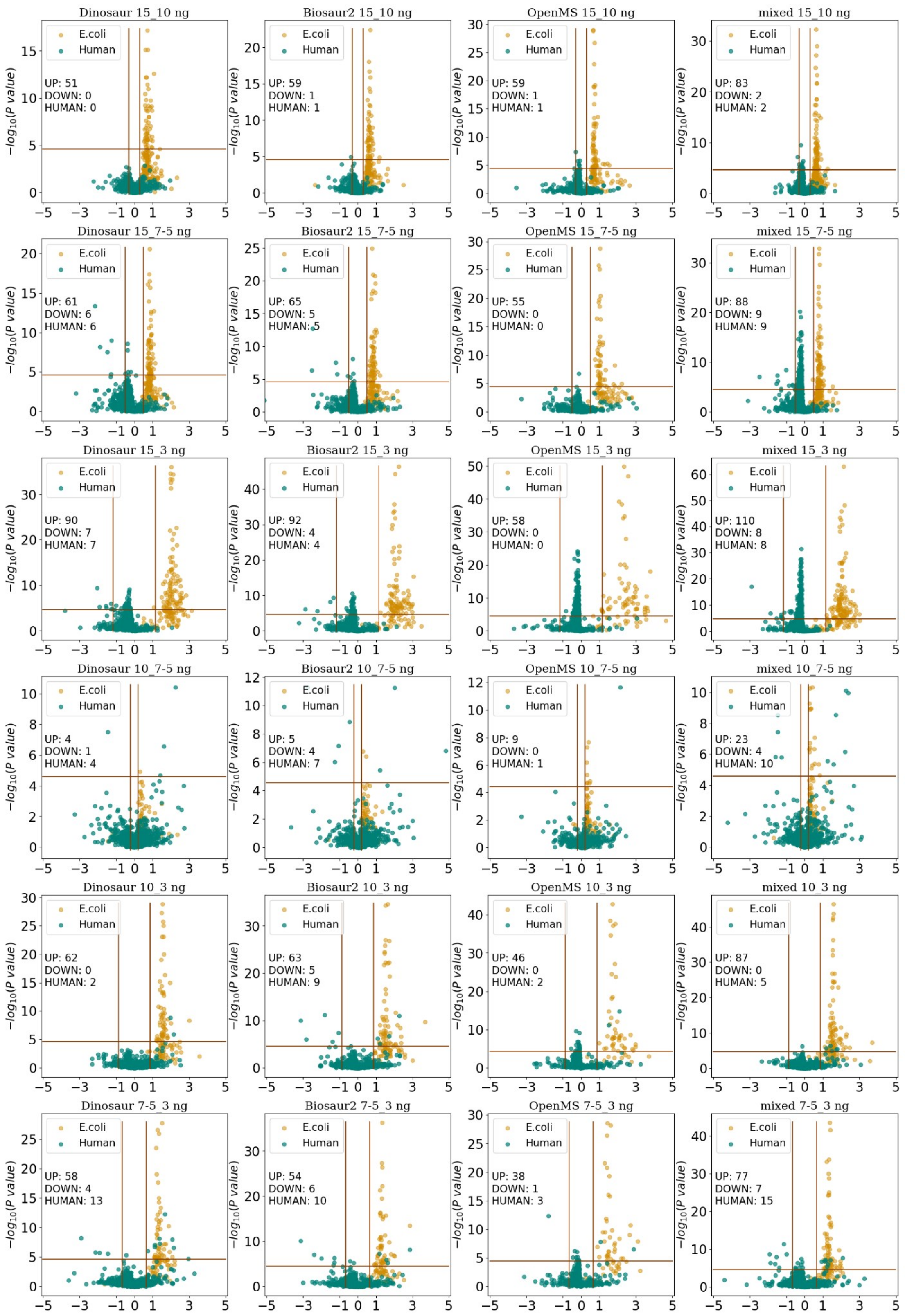


**Supplementary figure S3.** Volcano plots comparison for Human - *E. coli* dataset with turned on match-between-runs option. Horizontal line represents 0.05 p-value threshold adjusted using Bonferroni correction, vertical lines represent fold change thresholds which were set at a half of the real *E. coli* proteins ratio.

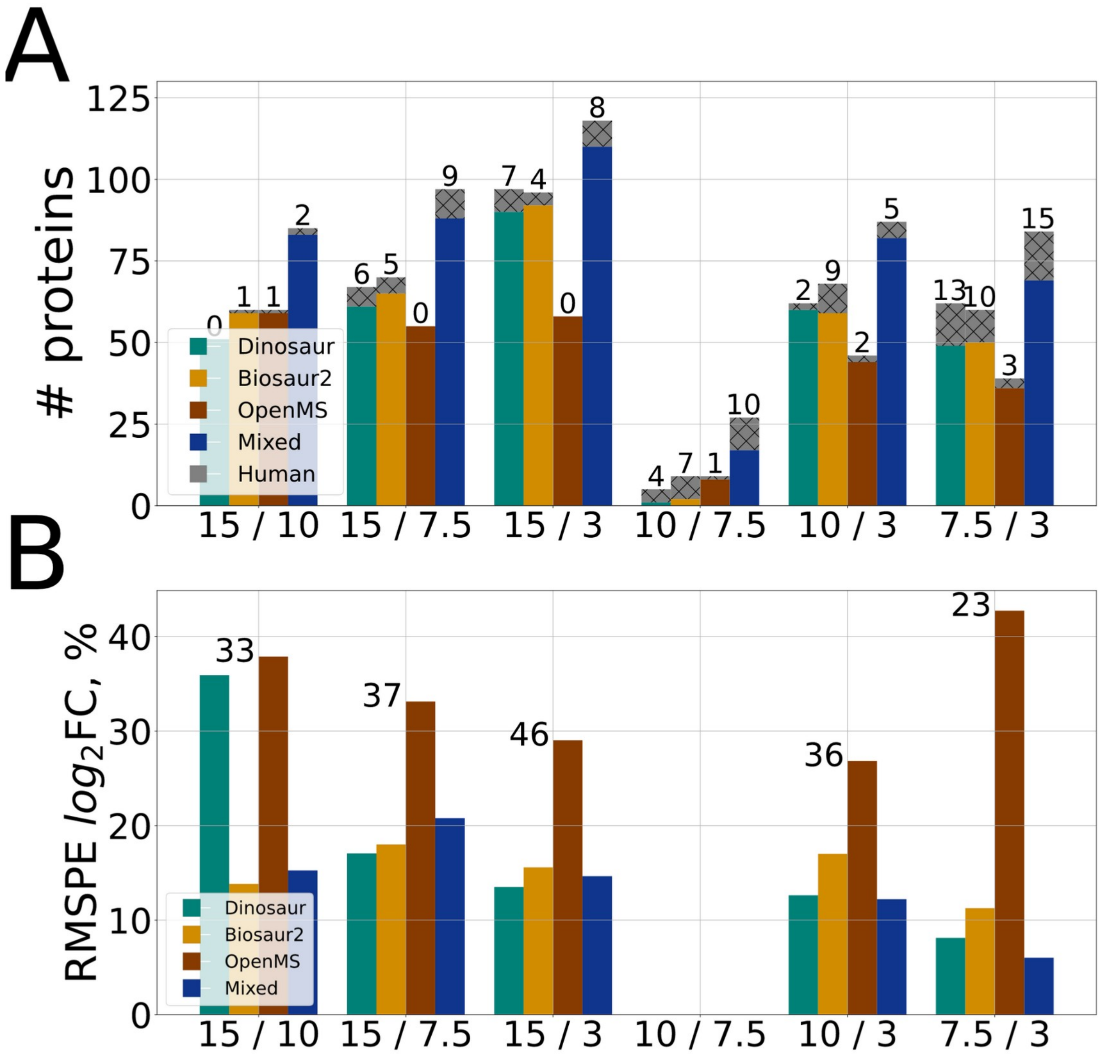


**Supplementary figure S4.** Histograms for six pairwise concentration comparisons for all HeLa - *E. coli* datasets using the IQMMA with turned on match-between-runs option. (A) the number of reported differentially expressed proteins. The numbers above the bars show the quantity of human proteins falsely identified as differentially expressed; (B) rooted mean squared percentage error of $\log_2$FC. The numbers above the bars show the size of the set of the common proteins for all four methods, for which the error was calculated
